# Left ventricular transcriptome and extracellular vesicle-derived miRNAs in porcine donation after circulatory death (DCD) hearts undergoing prolonged working mode *ex situ* perfusion

**DOI:** 10.1101/2025.01.22.634193

**Authors:** Fulin Wang, Eliana Lucchinetti, Phing-How Lou, Darren H. Freed, Michael Zaugg

**Author notes:** **<u>Corresponding Author</u>** Michael Zaugg, MD MBA FRCPC Department of Anesthesiology and Pain Medicine Department of Pharmacology University of Alberta CSB Room 2-150Y Edmonton AB T6G 2G3 (Canada). **Funding:** This study was funded by the Canadian Institutes of Health Research (CIHR), grant PJT173322 (MZ, DHF). **Competing Interests:** DHF is a member of the board of directors, Bridge to Life, Northbrook, IL 60062 (USA). All other authors declare no competing interests. **CRediT author statement**. **Fulin Wang:** data collection, data curation, formal analysis, writing - original draft preparation. **Eliana Lucchinetti:** conceptualization, data curation, formal analysis, writing - original draft preparation. **Phing-How Lou:** data collection. **Darren H. Freed:** conceptualization, methodology, data collection, funding acquisition. **Michael Zaugg:** conceptualization, project administration, writing - original draft preparation, funding acquisition.

## Abstract

Donation after circulatory death (DCD) hearts suffer from warm ischemia-reperfusion injury, which compromises graft quality. We have previously demonstrated that a postconditioning-based cardioprotective treatment (1% Intralipid, 2% (v/v) sevoflurane, 3 nM remifentanil) improved the function and viability of porcine DCD hearts undergoing *ex situ* heart perfusion (ESHP). However, transcriptional changes in DCD hearts subjected to prolonged working mode ESHP as compared with healthy hearts, and more so transcriptional changes in the presence or absence of cardioprotection, as well as the early reperfusion profile of extracellular vesicle (EV)-derived miRNAs, critical regulators of the transcriptional control, have not been investigated. A total of 5,483 differentially expressed transcripts were identified in left ventricular tissue samples of DCD hearts (N=10) collected after 6 hours of ESHP when compared with healthy hearts not subjected to ESHP (N=8), irrespective of cardioprotection. Downregulation of protein synthesis, metabolic, and DNA repair gene sets was most prominent, while inflammatory and remodeling gene sets were upregulated. Between DCD hearts that were treated with (pDCD_ESHP, N=5) or without (uDCD_ESHP, N=5) cardioprotection, only 43 transcripts were differentially regulated. Specifically, lipid breakdown, beta-oxidation, and DNA repair gene sets were upregulated in pDCD_ESHP, whereas lipid accumulation, inflammation, oxidative stress, and maladaptive remodeling gene sets were upregulated in uDCD_ESHP. Predominantly cardioprotective EV-derived miRNAs were released into the perfusate in pDCD_ESHP, while predominantly cardiac injury-associated miRNAs were released into the perfusate in uDCD_ESHP. Cardioprotection promoted adaptive rather than maladaptive transcriptional changes and enabled the release of potentially beneficial miRNAs.

## INTRODUCTION

Donation after circulatory death (DCD) hearts are a valuable source of donor organs, which can alleviate the ongoing shortage of suitable donor hearts. Recent studies have demonstrated non-inferior short-term survival rates in recipients receiving a DCD heart as compared to standard donation after brain death (DBD) hearts [1]. However, clinicians are still reluctant to allocate DCD hearts to patients of a higher priority status, i.e., with more unstable conditions. Indeed, despite the non-inferior short-term survival outcomes, recipients with a DCD heart transplant experience higher rates of primary graft dysfunction, which may be associated with potentially less favorable long-term survival and higher health care costs [2]. As an unavoidable component of the circulatory death process, the warm ischemic injury contributes to the activation of inflammatory responses and cell death signaling pathways as well as suppression of oxidative phosphorylation, creating a complex myriad of adverse events, which impairs post-transplant outcomes [3].

We have previously shown that a combined cardioprotective cocktail applied at the onset of reperfusion preserved the function and viability of the DCD heart, which otherwise declined significantly over 6 hours of *ex situ* heart perfusion (ESHP) [4]. In addition, the unprotected DCD heart experienced higher levels of mitochondrial damage and developed a dysfunctional metabolic phenotype, typically associated with heart failure, including impaired oxidative phosphorylation and accumulation of cardiotoxic triglycerides in the left ventricular tissue [5]. The molecular basis and mechanisms for the observed phenotypic differences between the unprotected and protected DCD hearts have yet to be explored, but given the distinct functional and metabolic outcomes, we hypothesized that the cardioprotective treatment would generate a different left ventricular transcriptome compared with the unprotected DCD hearts. In addition, as ESHP is a strong stimulus alone and known to induce cardiac remodeling [6], we hypothesized that DCD hearts undergoing prolonged ESHP would have a fundamentally different transcriptomic signature from healthy hearts not subjected to the DCD-ESHP process, irrespective of any cardioprotective treatment. Deciphering such differences will ultimately allow for the identification of molecular targets and the development of novel therapeutic strategies that can further improve DCD heart outcomes.

Recently, there is increasing interest in extracellular vesicles (EVs), which contain cargo such as proteins, DNA, and RNA acting as critical mediators of cardiac remodeling [7]. In particular, the importance of microRNAs (miRNAs) have been highlighted in a number of cardiac diseases such as heart failure, myocardial infarction [8], as well as in heart transplantation and allograft rejection [9]. Conversely, miRNAs have also been found to be important mediators of cardioprotection by pharmacological conditioning [10]. miRNAs are able to modulate cellular gene expression by silencing mRNA molecules via complementary base pair binding, thus mediating cardiac injury or cardioprotection [11]. For example, propofol-induced cardioprotection is partially mediated by the upregulation of miRNA-541 to target *HMGB1* (high mobility group protein B1) and suppress apoptosis and myocardial injury [12]. Given the distinct outcomes between unprotected and protected DCD hearts, we hypothesize that a differential profile of EV-derived miRNAs would be present in the circulation of the DCD hearts during ESHP, with predominantly cardioprotective miRNAs present in the protected group and cardiac injury-related miRNAs in the unprotected group.

## MATERIALS AND METHODS

### Surgical protocol and ESHP

All procedures in this study were approved by the University of Alberta Animal Care and Use Committee (Edmonton, Alberta, Canada) and carried out in accordance with the Canadian Council on Animal Care (Ottawa, Ontario, Canada) guidelines. The surgical procedure and ESHP protocol have been previously described in detail (Fig. 1) [4].

**Fig 1.**
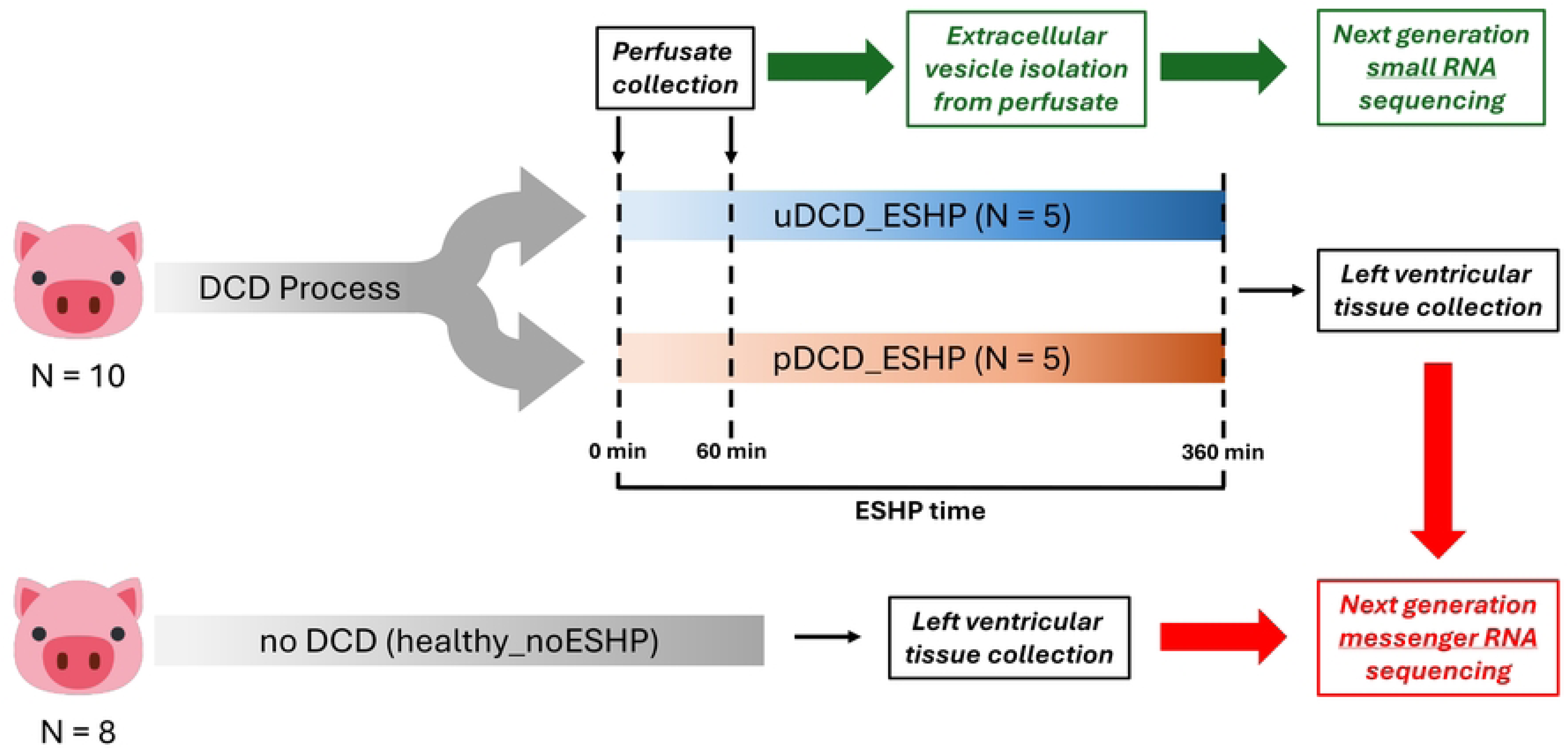
Experimental protocol. Anesthetized female pigs were subjected to a clinically identical DCD procedure after sternotomy. The DCD hearts were then mounted on an ESHP apparatus and perfused for a total of 6 hours. The hearts were randomly assigned to be perfused in the absence (uDCD_ESHP, N = 5) or presence (pDCD_ESHP, N = 5) of a cardioprotective pharmacological postconditioning treatment (1% Intralipid, 2% (v/v) sevoflurane, 3 nM remifentanil). Perfusate samples were collected before mounting of the heart (0 min) and after 60 min of ESHP time for profiling of extracellular vesicle-derived microRNAs. Left ventricular tissue was collected at the end of 6 hours of ESHP for profiling of tissue mRNAs. In addition, left ventricular tissue from hearts collected immediately after sternotomy served as healthy controls (healthy_noESHP, N = 8). DCD, donation after circulatory death; ESHP, *ex situ* heart perfusion. uDCD_ESHP, DCD hearts subjected to ESHP without cardioprotective treatment; pDCD_ESHP, DCD hearts subjected to ESHP with cardioprotective treatment; healthy_noESHP, hearts not subjected to the DCD process and ESHP.

Briefly, circulatory death was induced in female pigs (45.0 ± 4.5 kg) by the termination of mechanical ventilation followed by a stand-off period of 15 minutes prior to cardiectomy, mimicking the clinical protocol for DCD heart procurement. The DCD hearts were then rapidly excised and flushed with 1 L of tepid normokalemic adenosine-lidocaine cardioplegia solution before being mounted on a custom-made ESHP apparatus and perfused for a total of 6 hours. DCD hearts were randomly assigned to be perfused in the presence (protected, pDCD_ESHP, N=5) or absence (unprotected, uDCD_ESHP, N=5) of a multi-drug treatment consisting of Intralipid (1%), sevoflurane (2% v/v) and remifentanil (3 nM) applied at the onset of reperfusion. Intralipid remained in the circuit for the entire ESHP duration, while sevoflurane and remifentanil were only delivered to the circuit during the first 30 minutes of reperfusion [4]. DCD hearts were initially perfused in Langendorff mode (non-working) for 1 hour, and then switched to working mode for the remaining 5 hours. Cardiac tissue samples from the apex of the left ventricle were collected at the end of 6 hours of ESHP and rapidly snap-frozen in liquid nitrogen and stored at -80°C. Left ventricular tissue (apex) from hearts collected after sternotomy and not subjected to perfusion on the ESHP apparatus (healthy_noESHP, N=8) served as healthy controls. Perfusate samples were collected before mounting the hearts (baseline) on the ESHP apparatus and after one hour of ESHP (Fig. 1). The perfusate samples were centrifuged at 2’500g for 20 min at 4°C to separate plasma from cells. The plasma was passed through a 0.8 μm filter to remove debris and subsequently stored at -80°C.

### Next generation mRNA sequencing of LV

RNA extraction and library preparation were carried out on LV tissue samples using the mRNA-Seq (Poly-A) protocol at Fasteris SA (Plan-les-Ouates, Switzerland). Extracted RNA transcripts were purified by poly(A)-tail selection for mRNA. Libraries were constructed using the Illumina TruSeq Stranded mRNA Library Prep kit and sequenced on an Illumina Novaseq instrument. The libraries were mapped to the reference genome (Sscrofa11.1; https://www.ncbi.nlm.nih.gov/datasets/genome/GCF_000003025.6/) with the software package STAR. RSEM (RNA-Seq by Expectation-Maximization) was used for the quantification of RNA-seq data. Differential gene expression (DGE) analysis was carried out using the R package DESeq2, where genes with total counts under 10 were filtered out and normal transformation (log_2_[count + 1]) was applied to the filtered data [13]. The threshold was set to a minimum fold change of 1 and an adjusted p-value cutoff of 0.05. The RNASeq data was also subjected to gene set enrichment analysis (GSEA) [14, 15]. The gene sets used for the current analysis were obtained from the Molecular Signatures Database (MsigDB; https://www.gsea-msigdb.org/gsea/index.jsp), a collection of annotated gene sets for use with GSEA software, or were manually curated to include Bioplanet pathways (https://tripod.nih.gov/bioplanet/), WikiPathways (https://www.wikipathways.org/) or muscle-related pathways [16, 17]. The effects of the DCD-ESHP events were studied by combining pDCD_ESHP (N=5) and uDCD_ESHP (N=5) into the ex situ perfused DCD_ESHP phenotype (N=10) and then compared with healthy controls (healthy_noESHP) (N=8). The effects of the cardioprotective treatment were also investigated by comparing between pDCD_ESHP (N=5) and uDCD_ESHP (N=5). Gene sets were considered significantly enriched when the normalized enrichment score was greater than ± 1.35 and p-value < 0.1. DGE graphs were generated using R (version 4.4.2, R Foundation); volcano plots were generated using the EnhancedVolcano R package [18], heatmaps were generated using the pheatmap R package [19].

### Reverse transcription-quantitative real-time polymerase chain reaction (RT-qPCR)

Total RNA was extracted from frozen LV tissue using the Monarch® Total RNA Miniprep Kit (NEB, USA). The purified RNA was quantified and checked for purity (A_260/280_ ≥ 2.0 and A_260/230_ ≥ 1.8) using a NanoDrop 8000 Spectrophotometer (Thermo Fisher Scientific, USA). RT-qPCR reactions were set up using the Luna® Universal One-Step RT-qPCR Kit (NEB, USA) and performed in a LightCycler 480 (Roche, Switzerland). The primer information is listed in S1 Table. RT-qPCR data was analyzed using the LightCycler® 480 SW 1.5.1 (Roche, Switzerland) software. Ct (threshold cycle) values were determined using the second derivative method. Relative gene expression was computed using the ΔΔCt method, normalized to the healthy_noESHP group and using *ACTB* as the reference gene.

### EV isolation from perfusate

EVs were isolated from perfusate plasma samples by size exclusion chromatography (qEVoriginal, 70 nm isolation range, Izon Science, New Zealand) based on a previously published protocol [20]. Prior to loading onto the column, the perfusate plasma samples were additionally centrifuged at 1’500g for 10 min, followed by a second spin at 10’000g for 10 min at 4°C to remove any cellular debris and apoptotic bodies. 0.9 ml of clarified perfusate plasma was loaded onto the column (10 ml column volume, CV), which was pre-equilibrated with 1 CV of degassed PBS, pH 7.3. Eleven fractions of 0.5 ml were collected: fractions 1-4 consisted of the void volume; fractions 5-7 contained particles larger than the desired EV size range; fractions 8-10 were the fractions of interest for follow-up RNASeq and were enriched for EVs; fraction 11 was collected as a “post-elution” fraction in order to evaluate the presence of non-EV contaminants. The fractions were pooled into low-protein binding tubes and stored at -80°C. The columns were used for a maximum of 5 times and were cleaned with 2 CVs of degassed 0.1% Triton X-100 in PBS before re-equilibrating with 3 CVs of degassed PBS between uses.

### EV characterization by immunoblotting

Pooled EV fractions 1-4, 5-7, 8-10 and 11 were concentrated using 3 kDa MWCO Amicon® Ultra Centrifugal Filters (MilliporeSigma, USA) and solubilized using 4x RIPA buffer (NaCl [600 mM], Tris-HCl [200 mM], NP-40 [4% v/v], Triton X-100 [4% v/v], Sodium Deoxycholate [2% w/v], SDS [0.4% w/v], pH 8.0) containing a mixture of phosphatase and protease inhibitors. The mixture was vortexed and incubated on ice for 30 min to ensure total lysis. The protein content in the EV samples were quantified using the microBCA^TM^ Protein Assay Kit (Thermo Scientific, USA). EV samples were loaded at equal volume with the exception of fraction 11 (⅛ volume) due to its much higher protein content. The samples were separated by SDS-PAGE under reducing conditions and transferred onto 0.45 μm nitrocellulose membranes. The membranes were blocked using StartingBlock (TBS) Blocking Buffer (Thermo Scientific, USA), incubated with the appropriate primary and HRP-conjugated secondary antibodies, and developed using the ChemiDoc MP system (Bio-Rad Laboratories, USA). As per the Minimal Information for Studies of Extracellular Vesicles guidelines [21], a minimum of one protein from each category 1-3 were probed to evaluate EV purity: category 1a, CD81 (sc-166029, Santa Cruz, USA); category 2a, Tsg101 (GTX70255, GeneTex, USA), category 2b, cytoskeleton Vinculin (ab18058, Abcam, UK); category 3a, APOA1 (CAU30356, Biomatik, USA).

### Next generation sequencing of EV-derived miRNAs

RNA extraction and libraries were prepared using the small RNA gel free protocol at Fasteris, Life Science Genesupport SA (Plan-les-Ouates, Switzerland). Due to very low RNA yield in the baseline EV samples, all baseline samples from unprotected and protected groups were pooled. Libraries were constructed using the QIAseq miRNA library kit and sequenced on an Illumina NextSeq instrument. The readings were demultiplexed and mapped to the reference genome (Version Sscrofa11.1; https://www.ncbi.nlm.nih.gov/datasets/genome/GCF_000003025.6/) with the software package BWA. BEDTOOLS software was used for the quantification of RNASeq data. Differential gene expression (DGE) analysis was carried out using the R package DESeq2 [13] comparing between unprotected and protected hearts at 1 hour ESHP. Genes with total counts of less than 10 were filtered out, and shrinkage of effect size (log fold change estimates) was computed using the ashr R package [22].

### Evaluation of the degree of hemolysis in perfusate plasma

Perfusate plasma samples were processed with 2-step centrifugation to remove cellular debris and excess Intralipid (identical to sample preparation for EV isolation). The first spin was carried out at 1’500g for 10 min, followed by a second spin at 10’000g for 10 min at 4°C. The cleared plasma was diluted 10-fold in PBS and a spectra scan (350 - 750 nm) was performed in 96-well UV plates. The Hemolysis Score was calculated as previously described [23]. A threshold value of 0.057 was used to identify significantly hemolyzed samples. Difference in hemolysis score as a result of the treatment or time was compared using 2-way repeated measures ANOVA in R, followed by multiple comparison procedures (Bonferroni method) as appropriate. Correlation plots between hemolysis score and miRNA raw count abundance was generated using the ggplot2 R package [24] with linear regression R values and line equation annotated.

## RESULTS

Principal component analysis showed distinct transcriptomic phenotypes between the 3 groups, where the largest differences were observed between healthy controls (healthy_noESHP) and perfused DCD hearts (uDCD_ESHP and pDCD_ESHP), followed by a smaller, but still evident difference between unprotected (uDCD_ESHP) and protected (pDCD_ESHP) DCD hearts (Fig. 2A). Gene expression analysis identified a total of 5’483 differentially regulated transcripts between DCD hearts (DCD_ESHP) versus healthy controls (healthy_noESHP), of which 2’862 were upregulated and 2’621 were downregulated in DCD_ESHP hearts as compared to healthy_noESHP (Fig. 2B, S1 Figure). There was a small but significantly number of differentially regulated transcripts, i.e. 43, as a result of the cardioprotective treatment, whereby 21 were upregulated, and 22 were downregulated in the uDCD_ESHP group as compared to pDCD_ESHP (Fig. 2C and 2D).

**Fig. 2.**
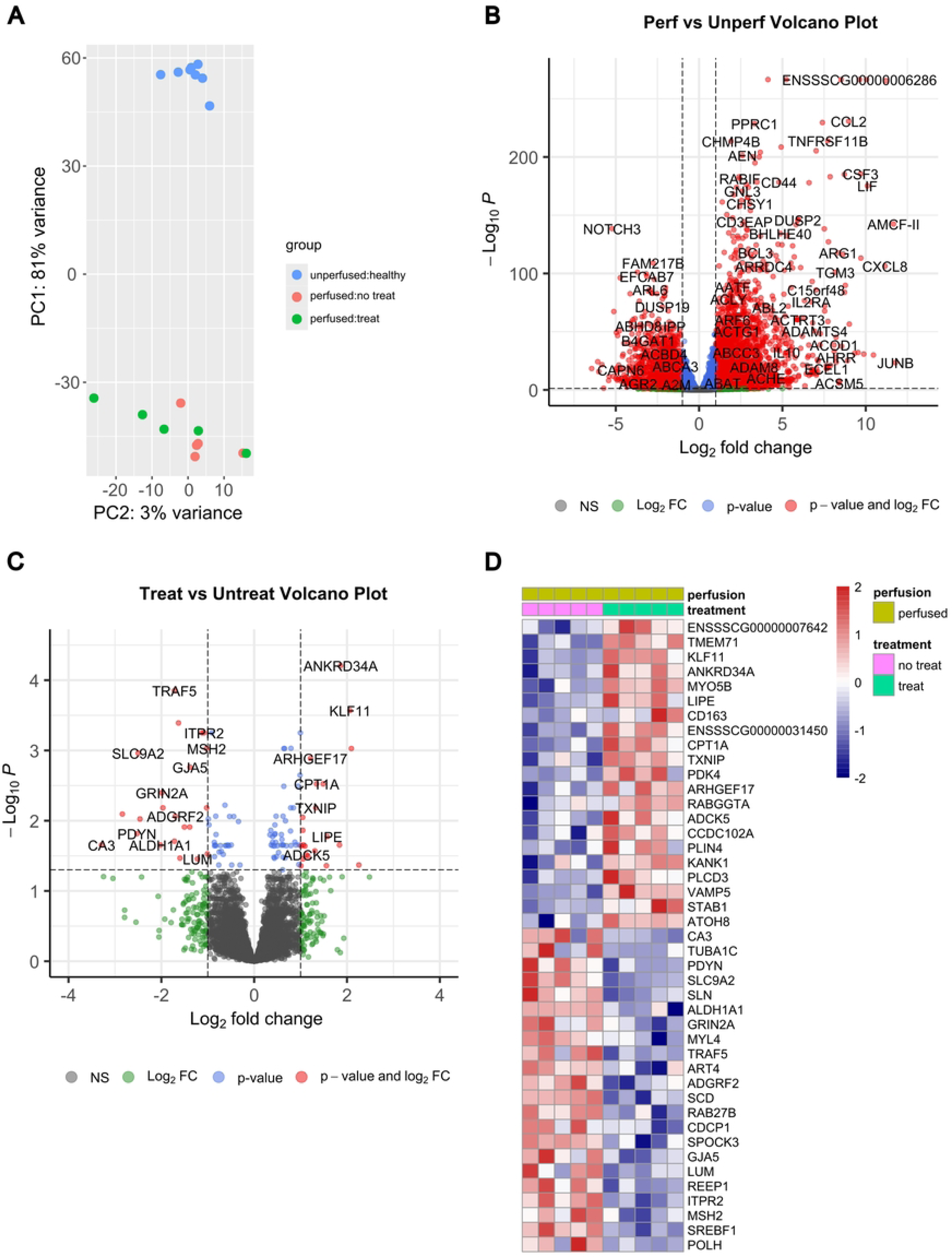
Overview of differential gene expression analysis in left ventricular tissue. <u>Panel A</u>. Unbiased principal component analysis plot of uDCD_ESHP, pDCD_ESHP, and healthy_ESHP groups. <u>Panel B</u>. Volcano plot of all detected mRNA transcripts from the comparison between perfused (uDCD_ESHP and pDCD_ESHP) and unperfused (healthy_ESHP) hearts. Differentially regulated genes (adjusted p-value < 0.05 and log_2_[fold change] > |± 1|) are highlighted in red. Top differentially regulated genes are annotated. <u>Panel C</u>. Volcano plot of all detected mRNA transcripts from the comparison between uDCD_ESHP and pDCD_ESHP hearts. Differentially regulated genes (adjusted p-value < 0.05 and log_2_[fold change] > ± 1 cutoff) are highlighted in red. Top differentially regulated genes are annotated. <u>Panel D</u>. Heatmap of all 43 differentially regulated genes between uDCD_ESHP and pDCD_ESHP groups. Gene expression is standardized to a mean of 0 and a variance of 1 (positive=upregulated, red; negative=downregulated, blue). DCD, donation after circulatory death; ESHP, *ex situ* heart perfusion; FC, fold change; PC, principal component; uDCD_ESHP, DCD hearts subjected to ESHP without cardioprotective treatment; pDCD_ESHP, DCD hearts subjected to ESHP with cardioprotective treatment; healthy_noESHP, hearts not subjected to the DCD process and ESHP.

### Transcriptional changes in DCD hearts undergoing prolonged ESHP compared with healthy controls not subjected to ESHP

*DCD-ESHP process downregulates fundamental regenerative and repair pathways* Transcript levels of mitochondrial tRNA synthetases, which load tRNAs with their respective amino acids for protein synthesis [25] (Fig. 3A), were reduced in DCD_ESHP hearts.

**Fig. 3.**
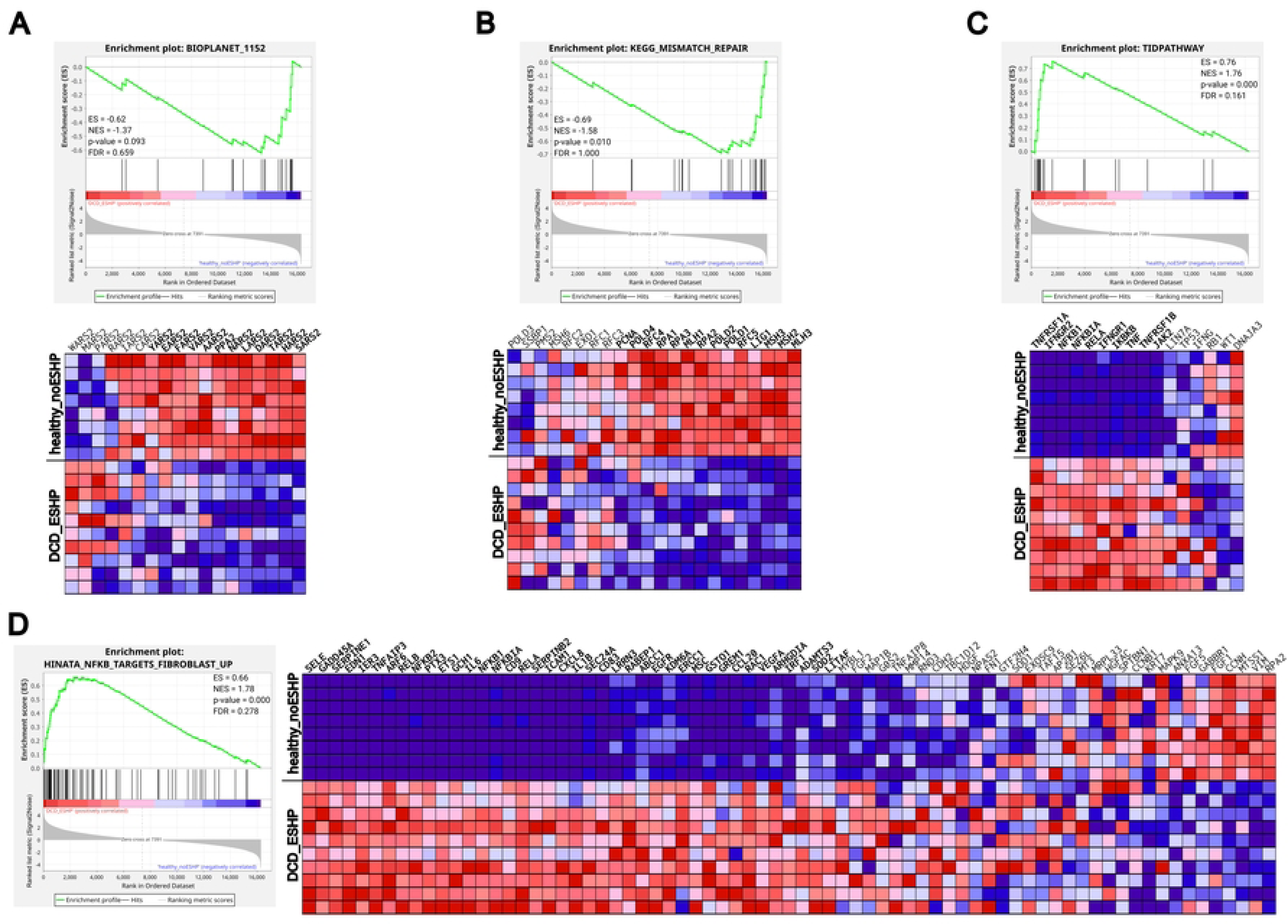
Gene set enrichment analysis comparing DCD hearts undergoing ESHP and healthy controls not subjected to ESHP. Enrichment plots and corresponding heatmaps of enriched gene sets. Enrichment plots are annotated with their enrichment score (ES), normalized enrichment score (NES), p-value and false discovery rate (FDR) for that particular gene set. Core enriched genes are in bold font in the heatmaps. <u>Panel A.</u> Mitochondrial tRNA aminoacylation (Bioplanet 1152). <u>Panel B</u>. KEGG (Kyoto Encyclopedia of Genes and Genomes) Mismatch Repair. <u>Panel C</u>. TID (also called DnaJ homolog subfamily A member 3 or DNAJA3) Pathway. <u>Panel D</u>. Hinata NFKB Targets Fibroblast Up (MsigDB; M4151). DCD, donation after circulatory death; ESHP, *ex situ* heart perfusion. DCD_ESHP, DCD hearts subjected to ESHP in the absence and presence of a cardioprotective treatment; healthy_noESHP, hearts not subjected to the DCD process and ESHP.

Conversely, the Arf pathway gene set, which inhibits ribosomal biogenesis by suppressing the transcription of rRNA (ribosomal RNA) [26], was enriched in DCD_ESHP hearts as compared to healthy_noESHP (S2 Table). Also, genes previously reported to be upregulated in C2C12 myotubes treated with cycloheximide, a potent inhibitor of protein translation, were enriched in DCD_ESHP [27]. In addition to protein synthesis gene sets, DNA strand elongation and replication, i.e. DNA synthesis, were downregulated in DCD_ESHP (S2 Table). In particular, DNA repair gene sets for single strand breaks, including base excision repair, nucleotide excision repair, mismatch repair, and double stranded breaks, as well as the telomerase reverse transcriptase gene set, responsible for telomere elongation, were downregulated in DCD_ESHP (Fig. 3B; S2 Table). Furthermore, metabolism-related gene sets such as NAD biosynthetic pathways and glycogen and fatty acid metabolism were downregulated in DCD_ESHP (S2 Table; S2 Figure, panel A and panel B).

#### DCD-ESHP process activates inflammatory pathways

Numerous inflammatory pathway gene sets were found to be enriched in DCD_ESHP. In particular, the Tid (tumorous imaginal disc) pathway [28] (Fig. 3C) and the LAIR (local acute inflammatory response) gene sets (S2 Figure, panel C), in addition to pro-inflammatory interleukin-2 (IL-2) and tumor necrosis factor (TNF) signaling pathway gene sets, were enriched in DCD_ESHP. Interestingly, the anti-inflammatory interleukin-10 (IL-10) signaling pathway gene set was also enriched in DCD_ESHP, likely as a counterregulatory response to the elevated inflammation. In addition, gene sets upregulated in response to pro-inflammatory toll-like receptor 4 (TLR4) agonist and lipopolysaccharide (LPS) treatments were similarly enriched in DCD_ESHP. Gene sets associated with immunity activation and chemotaxis [29–31] such as HCMV (human cytomegalovirus), CCR3 (C-C chemokine receptor type 3) and fMLP (N-formyl methionyl-leucyl-phenylalanine) pathways, were also enriched in DCD_ESHP (S2 Table).

#### DCD-ESHP process promotes remodeling processes

Several hypoxia-induced gene sets, including effector genes of hypoxia-inducible factor 1 (HIF1) were enriched in DCD_ESHP, likely due to the ischemic period during the circulatory death process. Gene sets associated with extracellular stress signaling such as transforming growth factor beta (TGFβ) (S2 Figure, panel D), vascular endothelial growth factor (VEGF) and endothelin-1 (ET-1) were also enriched in DCD_ESHP (S2 Table). Downstream p38, extracellular signal-regulated kinases (ERK), mitogen-activated protein kinases (MAPK), and NFκB signaling pathways were further upregulated in DCD_ESHP (S2 Figure, panel E; Fig. 3D). Remodeling of the LV in DCD_ESHP became specifically evident with enriched gene sets previously reported to be upregulated in transverse aortic constriction and differentiating C2C12 myotubes (S2 Table).

### Transcriptional changes in protected versus unprotected DCD hearts subjected to prolonged ESHP

#### Protected and unprotected DCD hearts have distinct metabolic signatures

Multiple gene sets associated with lipid accumulation were upregulated in uDCD_ESHP, including “SREBF and miR-33 in cholesterol and lipid homeostasis” (Fig. 4A) as well as “sphingolipid *de novo* biosynthesis” and fatty acid synthesis gene sets (S3 Table).

**Fig. 4.**
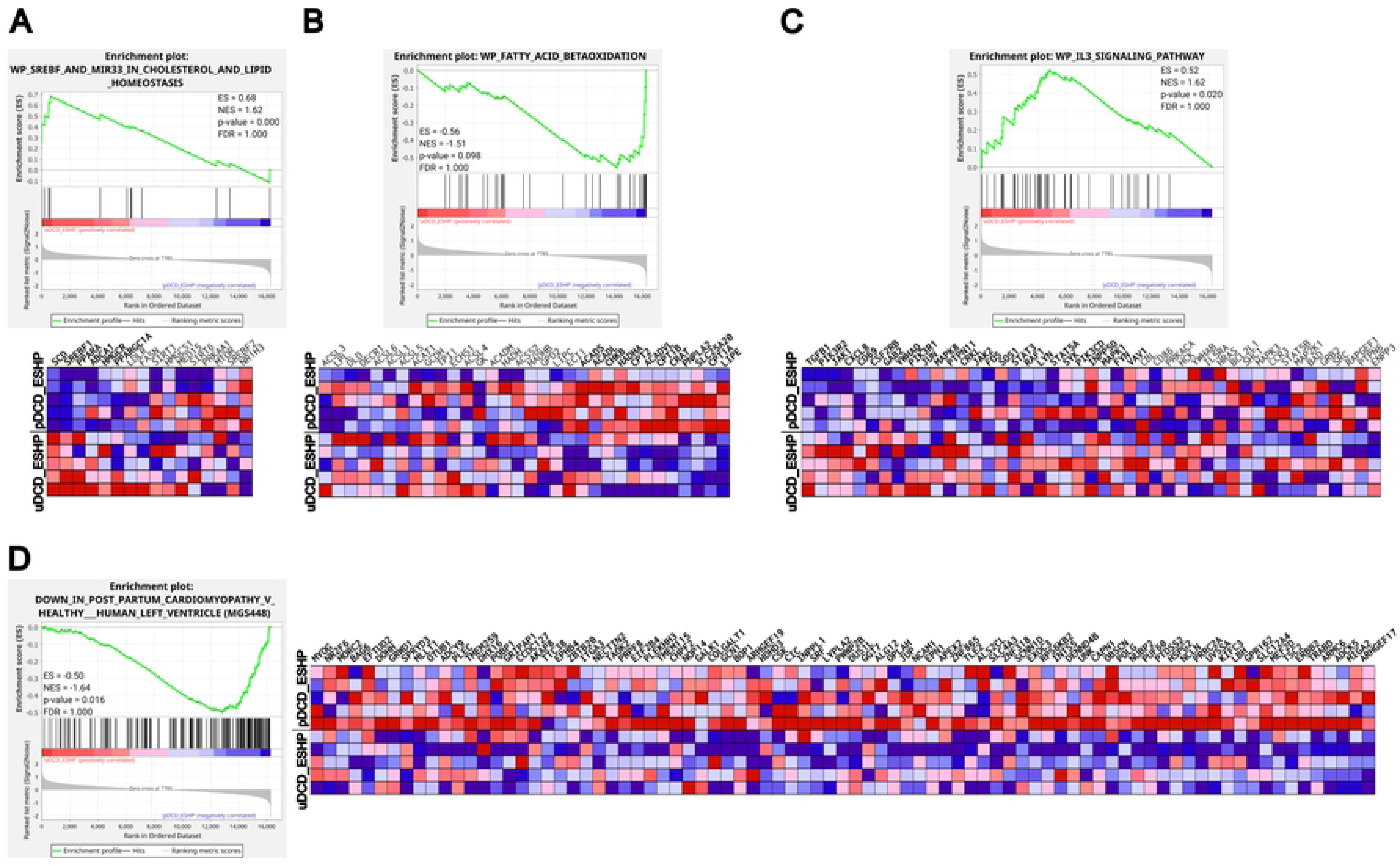
Gene set enrichment analysis comparing DCD hearts undergoing ESHP in the absence and presence of a cardioprotective treatment. Enrichment plots and corresponding heatmaps of enriched gene sets. Enrichment plots are annotated with their enrichment score (ES), normalized enrichment score (NES), p-value and false discovery rate (FDR) for that particular gene set. Core enriched genes are in bold font in the heatmaps. <u>Panel A.</u> SREBF and MiR33 in cholesterol and lipid homeostasis (WikiPathways WP2011). <u>Panel B</u>. Fatty acid beta oxidation (WikiPathways WP1269). <u>Panel C</u>. IL3 signaling pathway (WikiPathways WP286). <u>Panel D.</u> Down in postpartum cardiomyopathy vs. healthy human left ventricle (MuscleGeneSet MGS448); note the heatmap is cropped to only show the core enriched genes due to the large size of the gene set. DCD, donation after circulatory death; ESHP, *ex situ* heart perfusion. uDCD_ESHP, DCD hearts subjected to ESHP without cardioprotective treatment; pDCD_ESHP, DCD hearts subjected to ESHP with cardioprotective treatment; healthy_noESHP, hearts not subjected to the DCD process and ESHP.

Conversely, lipolysis and fatty acid oxidation gene sets were enriched in pDCD_ESHP (Fig. 4B; S3 Table). RT-qPCR analysis confirmed the differential expression of key regulatory genes involved in lipid metabolism in pDCD_ESHP vs uDCD_ESHP. Genes associated with lipid accumulation such as sterol response element binding factor 1 (*SREBF1*), peroxisome proliferator-activated receptor alpha (*PPARA*) and *GPAT4* glycerol-3-phosphate acyltransferase 4 (*GPAT4*) were significantly upregulated in uDCD_ESHP, while genes associated with fatty acid oxidation were significantly upregulated in pDCD_ESHP, including pyruvate dehydrogenase kinase 4 (*PDK4*), perilipin 2 (*PLIN2*), carnitine palmitoyltransferase 1A (*CPT1A*) and carnitine-acylcarnitine translocase (*SLC25A20*) (S4 Table). In addition, several amino acid metabolism gene sets (arginine/proline, alanine, histidine, tyrosine) as well as insulin signaling pathway were enriched in uDCD_ESHP, whereas the pentose phosphate pathway was enriched in pDCD_ESHP (S3 Table; S3 Figure, panel A).

#### Inflammation and oxidative stress pathways dominate in unprotected DCD hearts

Pro-inflammatory interleukin-2 (IL-2) and interleukin-6 (IL-6) pathways (S3 Figure, panels B-C), as well as the interleukin-3 (IL-3) signaling pathway (Fig. 4C) and the cytotoxic T-lymphocyte associated protein 4 (CTLA4) pathway (S3 Table) were upregulated in uDCD_ESHP. Core enriched genes not only included the cytokines themselves, but also their receptor subunits and downstream signaling effectors. TNF receptor associated factor 5 (*TRAF5*), a scaffold protein, which mediates TNF signaling [32], was significantly upregulated in uDCD_ESHP vs pDCD_ESHP (Fig. 2D). HIF1-responsive genes in the cardiovascular system were further upregulated in uDCD_ESHP, and were characterized by a signature similar to those exposed to oxidative stress (S3 Table). On the other hand, gene sets associated with DNA repair, particularly base excision repair and core enrichment of DNA glycosylases, were upregulated in pDCD_ESHP (S3 Figure, panel D). Furthermore, DNA polymerase eta (*POLH*), a low fidelity DNA polymerase which is activated in the presence of oxidative DNA damage [33], was significantly upregulated in uDCD_ESHP (Fig. 2D).

#### Maladaptive rather than adaptive remodeling occurs in unprotected DCD hearts

The dysregulated gene signatures of uDCD_ESHP resembled those in hearts with pressure overload or postpartum cardiomyopathies (Fig. 4D, S3 Table), both of which are characterized by LV dysfunction and reduced ejection fraction [34, 35]. The “induction of endoplasmic reticulum stress response by activating transcription factor 6 (ATF6)’’ gene set was upregulated in uDCD_ESHP, along with the core enriched gene DnaJ homolog subfamily C member 3 (*DNAJC3*), which is considered a protein marker for endoplasmic reticulum stress [36] (S3 Table). The p38- and ERK-MAPK pathway gene sets were enriched in uDCD_ESHP, including downstream effectors *FOS* (c-Fos), *JUN* (c-Jun), *MAPK8* (mitogen-activated protein kinase 8), subunits of NF-κB and signal transducer and activator of transcription 3 (*STAT3*). Subunits of NF-κB were core enriched in uDCD_ESHP, whereas NFκB inhibitor beta (*NFKBIB*), which suppresses the nuclear translocation of NFκB, was a core enriched gene in pDCD_ESHP (S3 Figure, panel E; S3 Table). The Spry pathway and Met pathway gene sets (S3 Figure, panel F; S3 Table), which interact with the MAPK signaling pathway [37, 38], were also enriched in uDCD_ESHP. Finally, the epidermal growth factor (EGF) and insulin-like growth factor 1 (IGF1) pathway gene sets, which converge onto MAPK signaling [39, 40], were upregulated in uDCD_ESHP (S3 Table).

### Distinct profiles of EV-bound miRNAs are present in the circulation of protected vs unprotected DCD hearts

Immunoblotting confirmed the presence of EV-positive CD81 and Tsg101 signals in F8-10, the EV-enriched fractions of interest, but not in the preceding F1-4 and 5-7. While the signal intensity of CD81 and Tsg101 was higher in F11, the levels of APOA1 and vinculin, a marker for non-EV contaminant particles [21], was also significantly higher in F11 and thus may have contained more cellular contaminants (S4 Figure). Therefore, F8-10 were used for the profiling of EV-derived miRNAs. DGE analysis identified 25 differentially regulated miRNAs between uDCD_ESHP and pDCD_ESHP with adjusted p-value <0.05 (Table 1). In addition, a literature search was conducted on the significantly regulated miRNAs to explore their previously reported functions in the cardiovascular system, specifically with regard to ischemia-reperfusion injury, as well as their putative targets and mechanisms of action. They were then broadly categorized into cardioprotective or cardiac injury-related miRNAs based on the reported findings (Table 1).

**Table 1.**
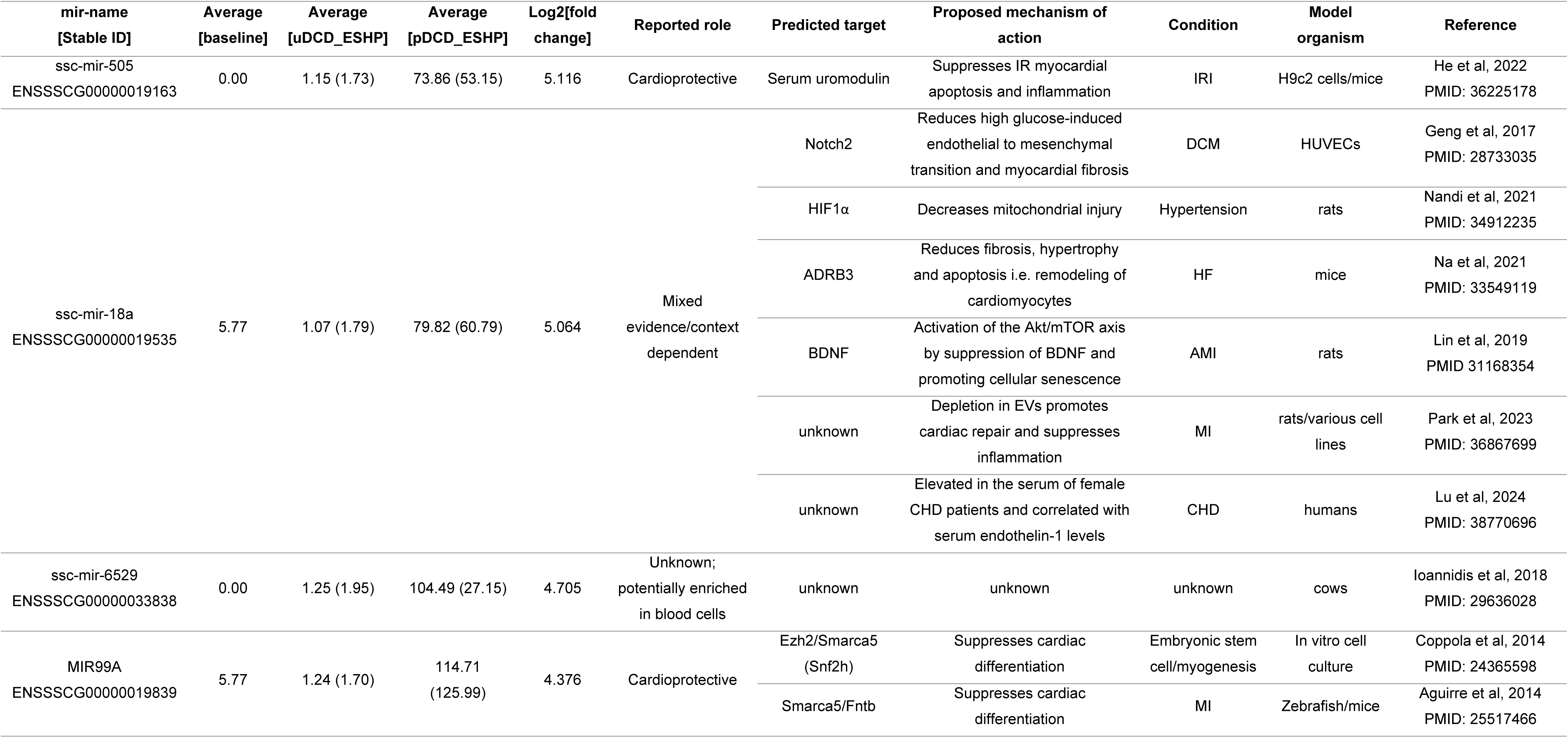
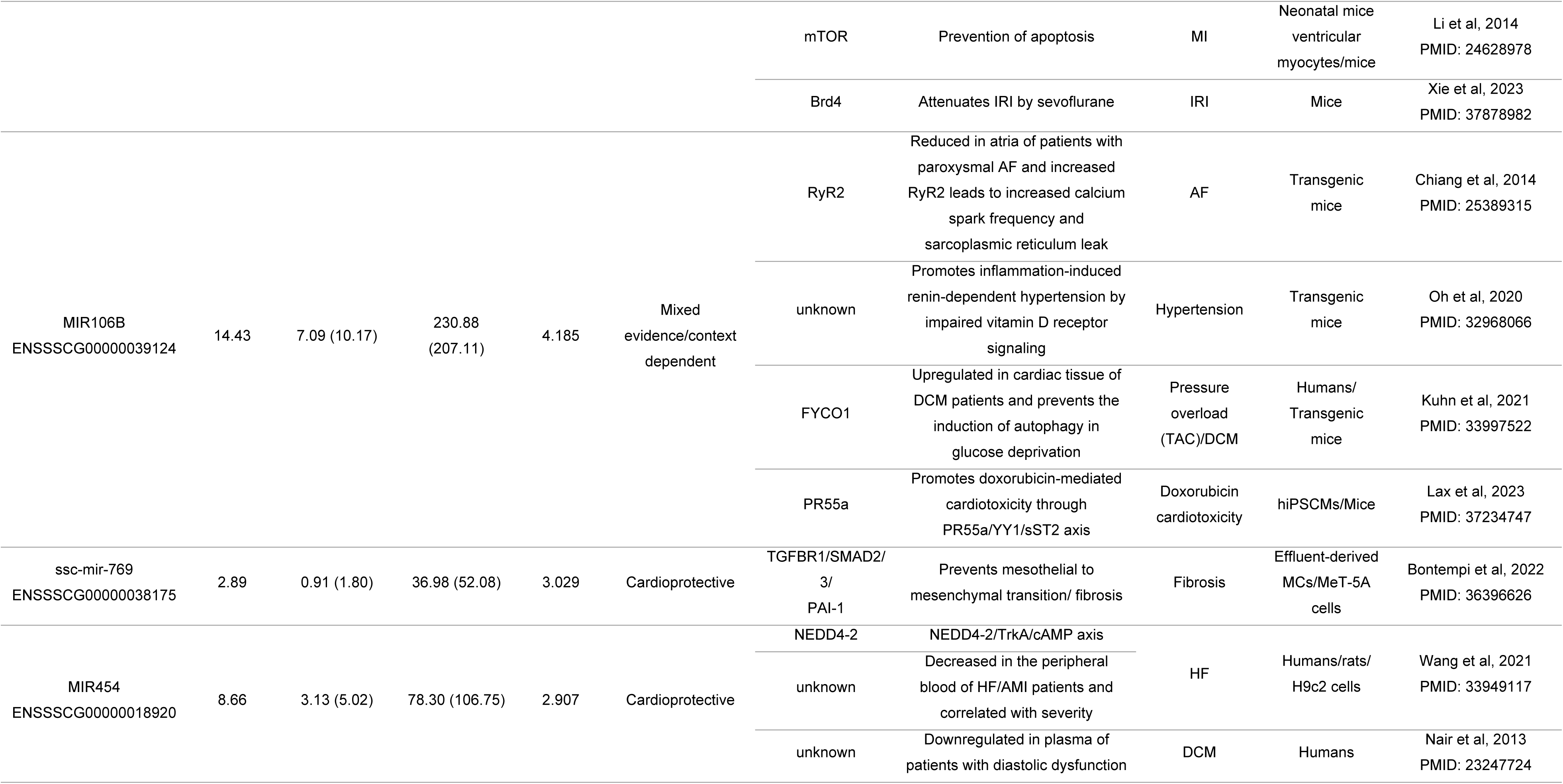
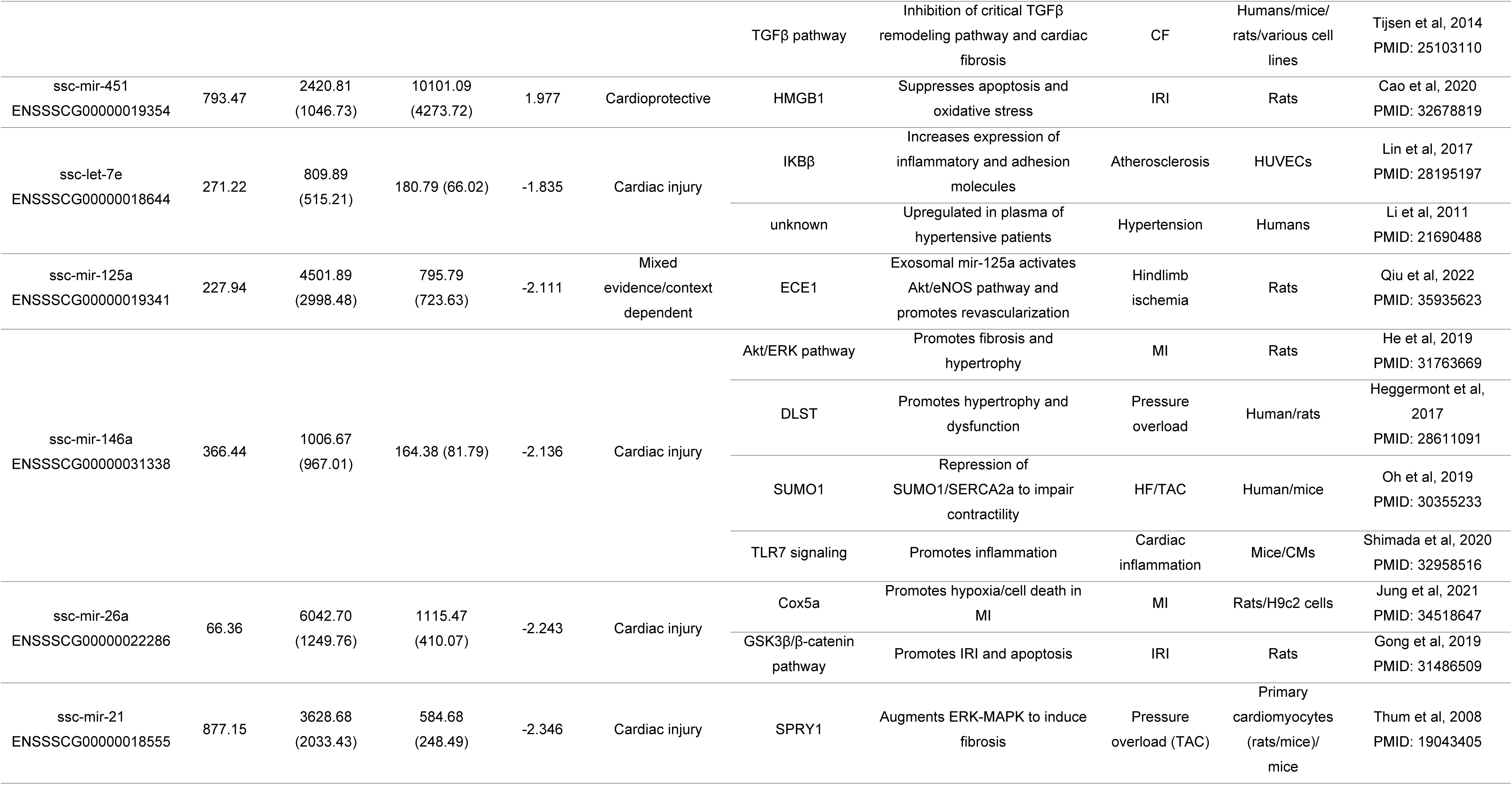
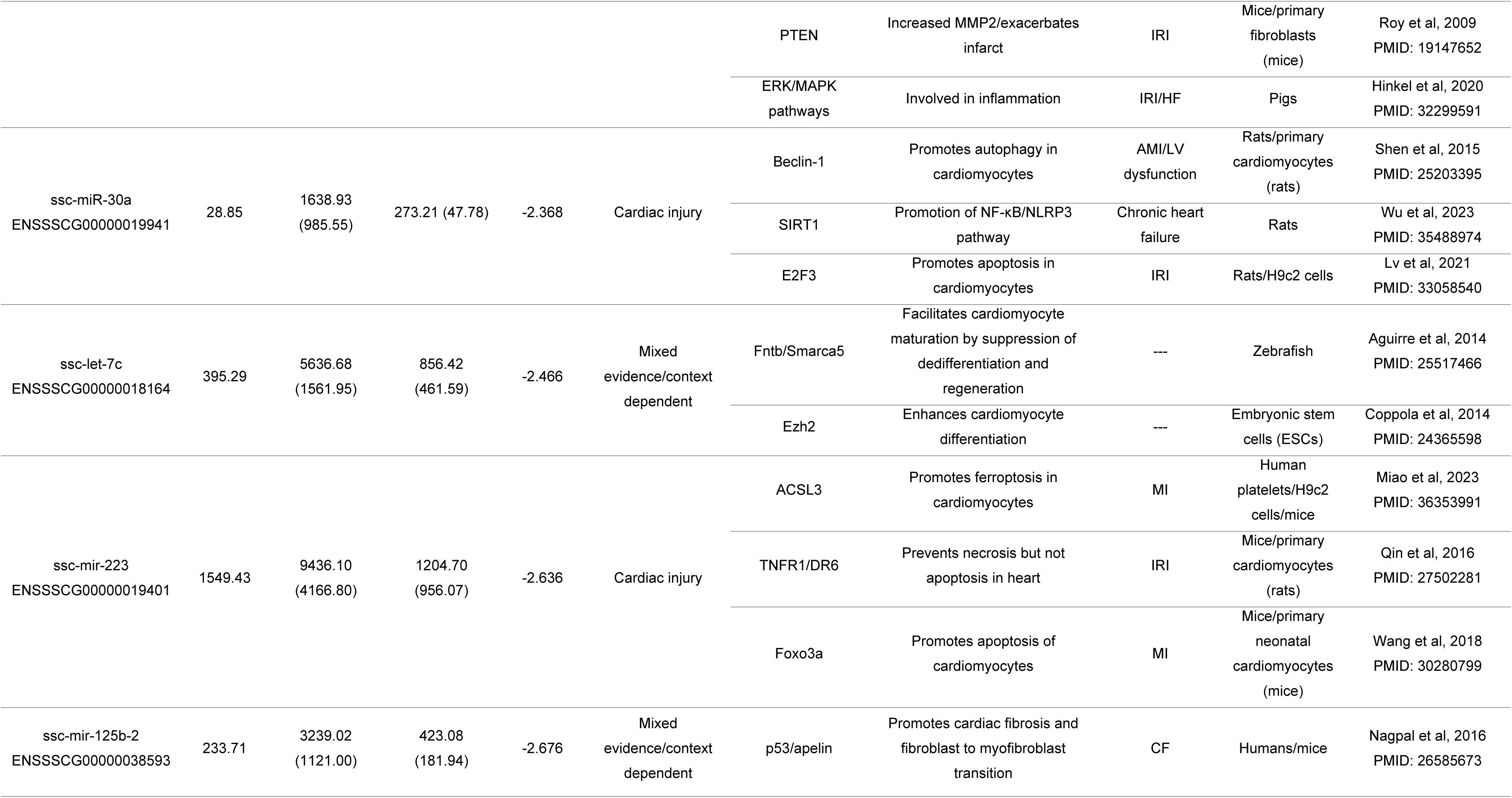
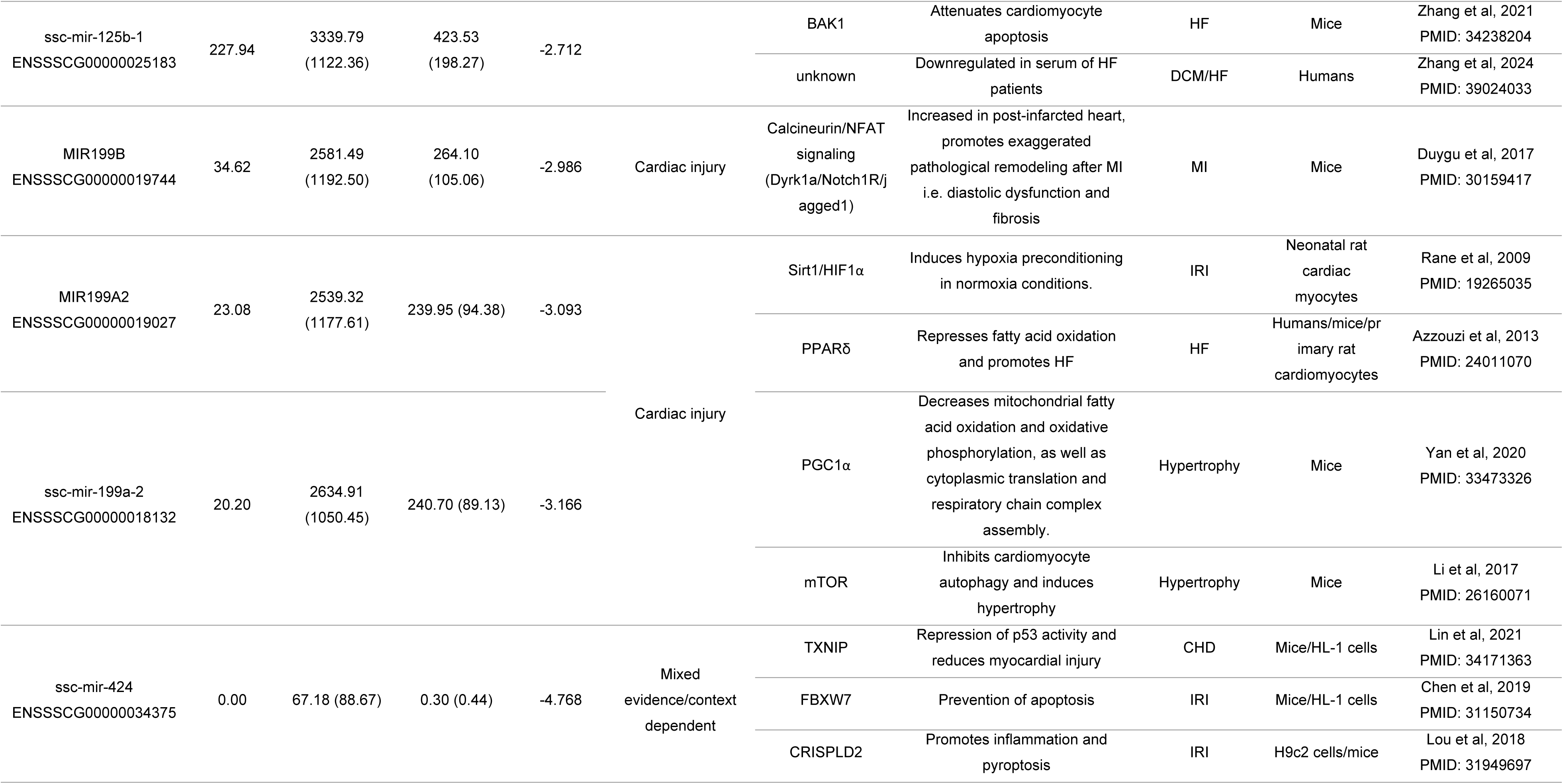
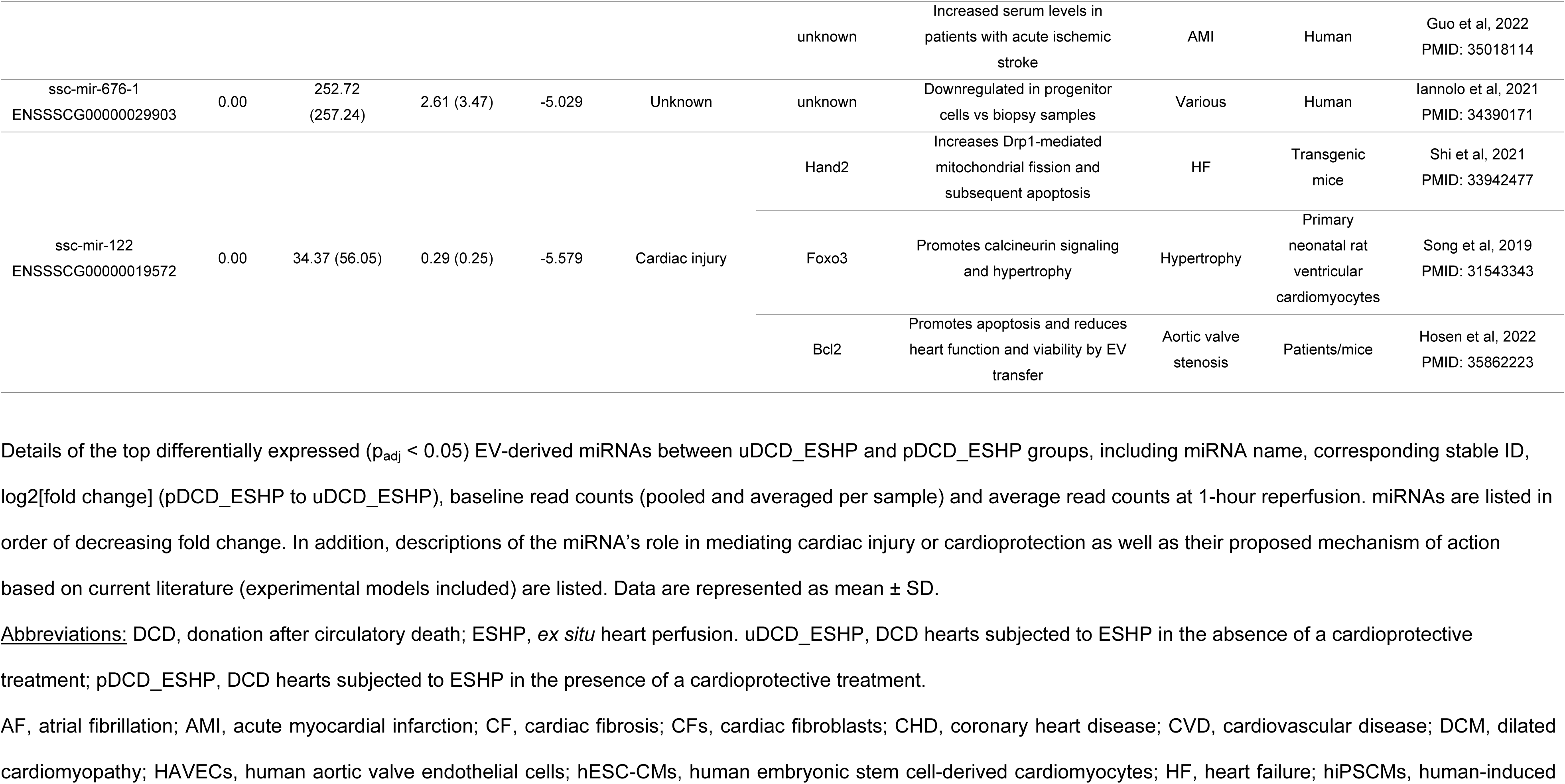
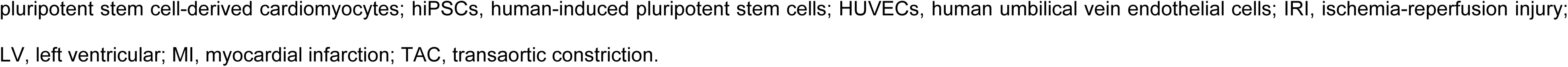
List of top differentially expressed extracellular-vesicle derived miRNAs at 1-hour reperfusion in uDCD_ESHP vs. pDCD_ESHP hearts.

miRNAs considered cardioprotective (ssc-mir-505, MIR99A, ssc-mir-769, ssc-mir-451, etc.) by preventing the activation of cell death signaling and differentiation of cells e.g. myofibroblasts, were released at higher levels from pDCD_ESHP compared to uDCD_ESHP. Conversely, miRNAs, considered cardiac injury-related (ssc-miR-30a, ssc-mir-26a, ssc-mir-21, ssc-mir-223, etc.) by promoting inflammation, fibrosis, and cell death, were released at higher levels from uDCD_ESHP. Some miRNAs have been reported to have ambiguous roles in cardiac injury versus cardioprotection, such as ssc-mir-424. In the myocardium, it can promote inflammation and pyroptosis by targeting cysteine-rich secretory protein LCCL domain-containing 2 (*CRISPLD2*), but also suppress apoptosis and p53 activity to reduce myocardial injury through F-box/WD repeat-containing protein 7 (*FBXW7*) and thioredoxin-interacting protein (*TXNIP*), respectively. In addition, elevated serum levels of this particular miRNA were observed in patients with acute ischemic stroke (Table 1).

Interestingly, certain erythrocyte-derived miRNAs [41], such as MIR106B, miR-486-2, miR-451, and miR-16, were present at higher levels in the protected group. These miRNAs also appear to be expressed at higher levels (p < 0.1) in pDCD_ESHP as compared to uDCD_ESHP (S5 Table). While perfusate hemolysis scores were not significantly different between pDCD_ESHP vs uDCD_ESHP (Fig. 5A), correlations of hemolysis scores with the erythrocyte-derived miRNAs showed R values ranging from 0.58 to 0.92, suggesting that hemolysis may have affected the miRNA profile (Fig. 5B).

**Fig. 5.**
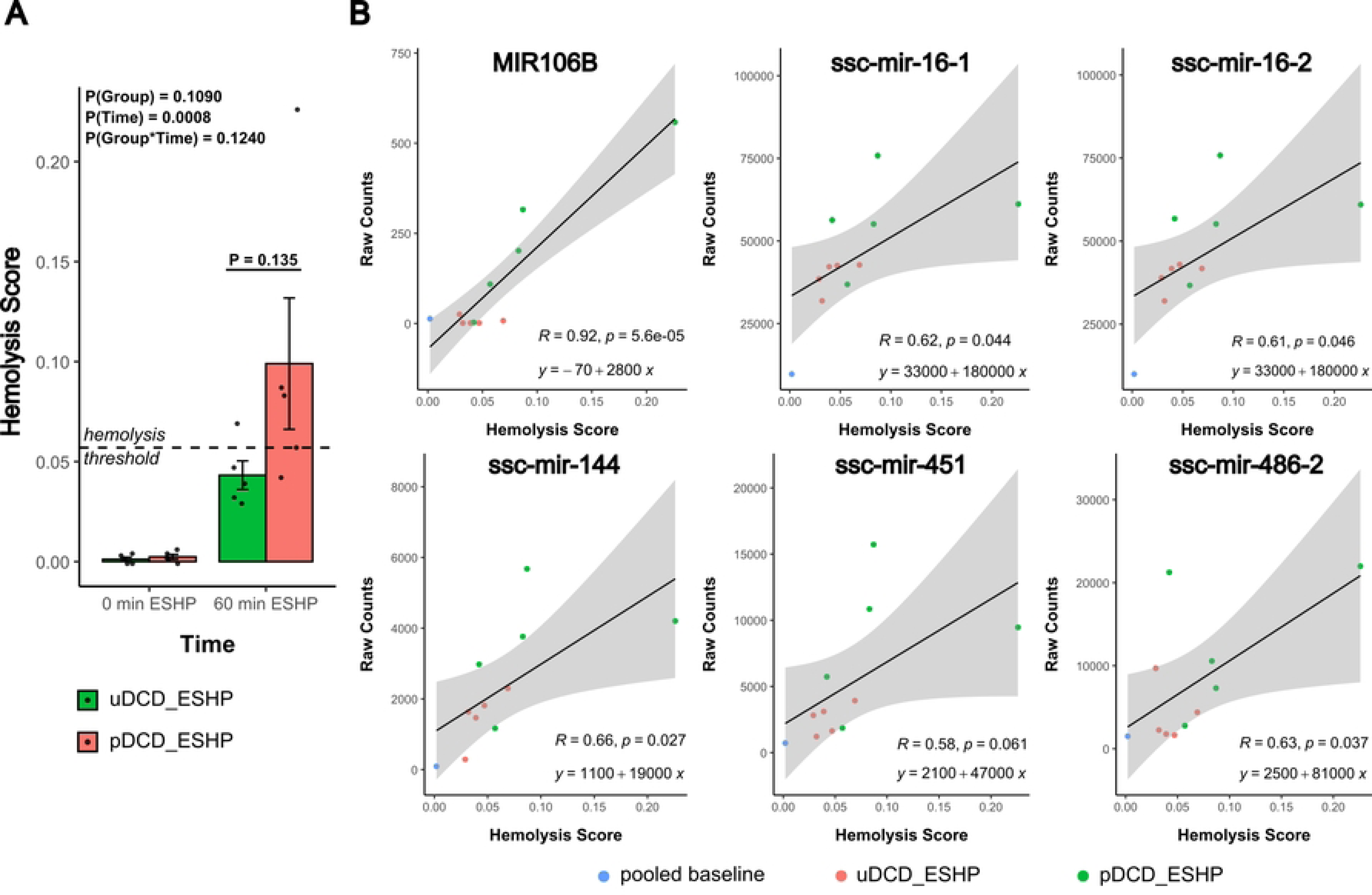
Degree of perfusate hemolysis and correlation with erythrocyte-related miRNAs in protected and unprotected DCD hearts. <u>Panel A.</u> Hemolysis score evaluation of perfusate samples of uDCD_ESHP and pDCD_ESHP hearts, collected at 0 min and at 60 min of ESHP. Measurements below the hemolysis score threshold of 0.057 (horizontal dashed line) are considered non-hemolyzed. Data is represented as mean ± SEM. <u>Panel B</u>. Correlation plots of hemolysis score with transcript counts of erythrocyte-related EV-derived miRNAs at 0 min and at 60 min ESHP. Linear correlation is expressed as R values and plotted as a dotted line. DCD, donation after circulatory death; ESHP, *ex situ* heart perfusion. uDCD_ESHP, DCD hearts subjected to ESHP without cardioprotective treatment; pDCD_ESHP, DCD hearts subjected to ESHP with cardioprotective treatment; healthy_noESHP, hearts not subjected to the DCD process and ESHP.

## DISCUSSION

Innovations such as the development of ESHP for donor heart preservation and evaluation contributed to the rapidly growing interest in including DCD hearts into clinical transplant programs. However, higher rates of primary graft dysfunction with the use of DCD hearts remain inadequately addressed [42, 43]. Previous studies predominantly focused on optimizing cardioplegia and ESHP perfusate compositions [44–46], and current clinical DCD heart transplant protocols only utilize limited strategies to improve the DCD heart graft quality. We have previously shown that ESHP provides a unique opportunity for donor heart protection by pharmacological postconditioning [4].

In an effort to better understand the injury process occurring in DCD hearts as well as putative repair mechanisms and pathways elicited by our cardioprotective treatment, a transcriptomics approach was chosen. With the focus on identifying potential novel therapeutic targets, the transcriptional changes as a result of the DCD-ESHP process were evaluated separately from the cardioprotective treatment effect, as this process *per se* was expected to evoke major transcriptional responses. Our study confirms that the DCD-ESHP process leads to the activation of many inflammation-related genes and pathways and further results in broad impairment of key energy-dependent processes such as RNA biogenesis and processing. Similar transcriptional changes have been reported in previous studies using cold storage of donor hearts [3] or other custom perfusion systems [47]. This suggests that detrimental transcriptional remodeling is inherent to many different types of cardiac preservation. However, our in-depth analysis identified many additional transcriptional changes in DCD hearts, which can be used to develop new therapeutic strategies.

Energy substrate metabolism is central in the development of cardiomyopathies and has been explored as a therapeutic target for quite some time. Unprotected DCD hearts exhibited severe dysregulation of lipid metabolism and increased expression of lipid synthesis genes e.g. *SCD, SREBF1*, a phenotype which is often seen in heart failure [48]. In protected DCD hearts, *PDK4* was significantly upregulated in accordance with previously confirmed increased fatty acid oxidation [4], while utilization of glucose as energy substrate is expected to be suppressed [49]. Rather than entering the Krebs cycle, this excess glucose may likely be shuttled into the pentose phosphate cycle instead, contributing to the maintenance of a healthy redox balance and nucleic acid synthesis in protected hearts. Interestingly, in unprotected DCD hearts, a number of amino acid metabolism gene sets were enriched, suggesting that amino acids may serve as “rescue” energy substrates, specifically in the ischemic heart with its limited oxygen supply and impaired fatty acid oxidation [50]. Considering that protected and unprotected DCD hearts displayed distinct metabolic phenotypes, targeting metabolic fuel preference appears to be a highly promising strategy to further improve DCD heart outcomes.

Preventing oxidative damage, a classic hallmark of myocardial ischemia-reperfusion injury, may be another promising target in improving DCD heart outcomes. As DNA repair gene sets were enriched in protected DCD hearts, the decreased ability of unprotected DCD hearts to excise and remove DNA lesions, such as 8-oxoguanine, may have led to the induction of *POLH* (DNA polymerase eta), a low fidelity DNA polymerase which is activated in the presence of DNA damage [51]. In addition, base excision repair gene sets, the major DNA repair mechanism in mitochondria, were upregulated in protected DCD hearts, whereas the gene response to oxidative stress was upregulated in unprotected DCD hearts, a possible sign of more severe injury. As more therapies targeting DNA damage repair are being explored in the context of various heart diseases such as heart failure and myocardial infarction, this strategy may also prove to be useful in DCD heart transplants [52]. Preventing oxidative damage to proteins would further reduce the need for protein turnover, namely degradation of the damaged proteins and *de novo* synthesis, an energy consuming process in the cell. Since oxidative phosphorylation and thus ATP production is diminished in unprotected DCD hearts [4], ATP-consuming aminoacyl-tRNA synthetases and chaperone proteins, which participate in protein synthesis and folding of *de novo* and damaged proteins, are likely to be more impaired in unprotected DCD hearts and thus can only contribute to a limited recovery. Transcripts associated with protein synthesis capacity were equally diminished in both untreated and treated DCD hearts, as commonly seen in myocardial ischemia reperfusion injury [53]. However, unprotected DCD hearts had higher levels of oxidized proteins [4], and accordingly the endoplasmic reticulum stress response gene set [54] was enriched in these hearts.

Emerging therapeutic strategies have begun to target inflammatory pathways and signaling cascades (e.g. NF-κB and MAPK) in the treatment of heart failure and cardiovascular diseases [55]. In unprotected DCD hearts, pro-inflammatory IL-2 and IL-6 pathway gene sets were upregulated. In addition, ERK-MAPK signaling gene sets were enriched in unprotected DCD hearts, including the core enriched gene *MAPK8*, which is associated with cell death and may exacerbate cardiac injury [56]. Interestingly, the combination of impaired DNA repair, enhanced pro-inflammatory cytokines, and ERK-MAPK signaling pathways in unprotected DCD hearts recapitulates the key features of the senescence-associated secretory phenotype (SASP) of the heart, which contributes to the development of atherosclerosis, hypertension, as well as increased morbidity and mortality [57]. Thus, a key strategy for improving outcomes in DCD transplants may lay in the suppression of cell death pathways and targeting remodeling processes in the heart.

Many identified miRNAs in the perfusate of unprotected DCD hearts were linked to the c-Myc, NF-κB and MAPK signaling pathways and may exert their deleterious effects by promoting differentiation and cell death pathways in the heart. As the significance of miRNAs in the field of cardiovascular and heart transplant research has become increasingly important in recent years [9, 58], our novel findings related to EV-derived miRNAs in the perfusate of DCD hearts during ESHP open up an exciting new avenue of studying their mechanistic role in DCD heart preservation and their potential as biomarkers as well as therapeutic targets. It is important to mention that in our setting the miRNAs may be released from sources other than the heart, suggesting that the cardioprotective treatment was able to condition the release of potentially beneficial miRNAs from blood cells present in the perfusate. Indeed, many of the abundant miRNAs observed in this study, such as miR-125, miR-223 and let-7, have also been reported to be highly expressed in blood cells, including erythrocytes [41], platelets [59], and macrophages [60]. In particular, the activation of platelets in response to stimuli such as inflammation and vascular injury can release EVs [61], which contain miRNAs and participate in intercellular communication. Interestingly, many of the platelet-associated miRNAs were upregulated in uDCD_ESHP, which suggests a higher degree of platelet activation and inflammation. Activated platelets release mitochondria as a damage-associated molecular pattern [62], and the abundance of cell-free mitochondrial DNA was indeed higher in the uDCD_ESHP group [4]. Clinical and experimental evidence shows that sevoflurane specifically attenuates inflammatory processes in platelets [63, 64], further supporting the use of our DCD organ protective strategy during ESHP.

We would like to mention the following limitations of our study. While profiling of the transcriptome provides a broad overview of the cellular landscape, the biological actions of the transcripts are ultimately carried out at the protein level. Crucial information including protein expression, post-translational modifications and protein-protein interactions could not be assessed in this transcriptomics study and should be explored in follow-up studies. In addition, only LV tissue mRNAs and perfusate EV-derived miRNAs were investigated in this study. Future studies should also investigate transcriptional changes in other parts of the DCD heart, such as the right ventricle and coronary arteries, as well as incorporate the evaluation of time-lapse transcriptional changes. As hemolysis occurred in the perfusate samples as a result of the ESHP process, the presence of erythrocyte-derived miRNAs interfered with the profiling of EV-associated miRNAs and thus should be accounted for (e.g. correlation of hemolysis score and abundance of blood-related miRNAs) and interpreted with caution [65]. Since the perfusate used in this study contained autologous blood, the perfusate composition should be evaluated (e.g. erythrocyte-free buffers) depending on the objective. In addition, the predicted targets of miRNAs have been predominantly studied in non-porcine models and should be validated using the porcine genome. Finally, while juvenile pigs were chosen for this study in order to minimize the effects of sex hormones, there have been reported sex differences even in juvenile animals [66, 67].

### Conclusions

The DCD-ESHP process profoundly reduced transcripts associated with the metabolic and protein synthesis capacity and increased transcripts involved in inflammation and remodeling of the left ventricle. A detailed comparison of the left ventricular transcriptome of protected and unprotected DCD hearts unveiled energy substrate metabolism, redox balance with its implications on DNA repair and protein homeostasis, and inflammation-associated cell death signaling as the most promising therapeutic targets for improving outcomes in DCD heart transplants. The identification of differentially expressed EV-derived miRNAs in the perfusates of protected versus unprotected DCD hearts opens a completely new avenue of research.

### Data availability statement

The raw mRNA and miRNA sequence data for this paper is available at NCBI BioProject repository under the accession number PRJNA1002377.

## REFERENCES

1. Schroder JN, Patel CB, DeVore AD, Bryner BS, Casalinova S, Shah A, et al. Transplantation Outcomes with Donor Hearts after Circulatory Death. N Engl J Med. 2023;388(23):2121–31. doi: 10.1056/NEJMoa2212438.

2. Squiers JJ, DiMaio JM, Van Zyl J, Lima B, Gonzalez-Stawisnksi G, Rafael AE, Meyer DM, Hall SA. Long-term outcomes of patients with primary graft dysfunction after cardiac transplantation. Eur J Cardiothorac Surg. 2021;60(5):1178–83. doi: 10.1093/ejcts/ezab177.

3. Lei I, Wang Z, Chen YE, Ma PX, Huang W, Kim E, et al. "The Secret Life of Human Donor Hearts": An Examination of Transcriptomic Events During Cold Storage. Circ Heart Fail. 2020;13(4):e006409. doi: 10.1161/CIRCHEARTFAILURE.119.006409.

4. Wang F, Lucchinetti E, Lou PH, Hatami S, Chakravarty A, Hersberger M, Freed DH, Zaugg M. Optimizing resuscitation of the donation after circulatory death heart by mitochondrial protection in a female porcine model. Anesthesiology. 2024. doi: 10.1097/ALN.0000000000005093.

5. Bertero E, Maack C. Metabolic remodelling in heart failure. Nat Rev Cardiol. 2018;15(8):457–70. doi: 10.1038/s41569-018-0044-6.

6. Movahed M, Brockie S, Hong J, Fehlings MG. Transcriptomic Hallmarks of Ischemia-Reperfusion Injury. Cells. 2021;10(7). doi: 10.3390/cells10071838.

7. Patil M, Henderson J, Luong H, Annamalai D, Sreejit G, Krishnamurthy P. The Art of Intercellular Wireless Communications: Exosomes in Heart Disease and Therapy. Front Cell Dev Biol. 2019;7:315. doi: 10.3389/fcell.2019.00315.

8. Colpaert RMW, Calore M. MicroRNAs in Cardiac Diseases. Cells. 2019;8(7). doi: 10.3390/cells8070737.

9. Shah P, Agbor-Enoh S, Bagchi P, deFilippi CR, Mercado A, Diao G, et al. Circulating microRNAs in cellular and antibody-mediated heart transplant rejection. J Heart Lung Transplant. 2022;41(10):1401–13. doi: 10.1016/j.healun.2022.06.019.

10. Roth S, Torregroza C, Feige K, Preckel B, Hollmann MW, Weber NC, Huhn R. Pharmacological Conditioning of the Heart: An Update on Experimental Developments and Clinical Implications. Int J Mol Sci. 2021;22(5). doi: 10.3390/ijms22052519.

11. Zhou SS, Jin JP, Wang JQ, Zhang ZG, Freedman JH, Zheng Y, Cai L. miRNAS in cardiovascular diseases: potential biomarkers, therapeutic targets and challenges. Acta Pharmacol Sin. 2018;39(7):1073–84. doi: 10.1038/aps.2018.30.

12. Li YM, Sun JG, Hu LH, Ma XC, Zhou G, Huang XZ. Propofol-mediated cardioprotection dependent of microRNA-451/HMGB1 against myocardial ischemia-reperfusion injury. J Cell Physiol. 2019;234(12):23289–301. doi: 10.1002/jcp.28897.

13. Love MI, Huber W, Anders S. Moderated estimation of fold change and dispersion for RNA-seq data with DESeq2. Genome Biol. 2014;15(12):550. doi: 10.1186/s13059-014-0550-8.

14. Mootha VK, Lindgren CM, Eriksson KF, Subramanian A, Sihag S, Lehar J, et al. PGC-1alpha-responsive genes involved in oxidative phosphorylation are coordinately downregulated in human diabetes. Nat Genet. 2003;34(3):267–73. doi: 10.1038/ng1180.

15. Subramanian A, Tamayo P, Mootha VK, Mukherjee S, Ebert BL, Gillette MA, et al. Gene set enrichment analysis: a knowledge-based approach for interpreting genome-wide expression profiles. Proc Natl Acad Sci U S A. 2005;102(43):15545–50. doi: 10.1073/pnas.0506580102.

16. Lucchinetti E, Hofer C, Bestmann L, Hersberger M, Feng J, Zhu M, et al. Gene regulatory control of myocardial energy metabolism predicts postoperative cardiac function in patients undergoing off-pump coronary artery bypass graft surgery: inhalational versus intravenous anesthetics. Anesthesiology. 2007;106(3):444–57. doi: 10.1097/00000542-200703000-00008.

17. Malatras A, Duguez S, Duddy W. Muscle Gene Sets: a versatile methodological aid to functional genomics in the neuromuscular field. Skelet Muscle. 2019;9(1):10. doi: 10.1186/s13395-019-0196-z.

18. Blighe K, Rana S, Lewis M. EnhancedVolcano: publication-ready volcano plots with enhanced colouring and labeling [Internet]. 2021; Available from: https://github.com/kevinblighe/EnhancedVolcano.

19. Kolde R. pheatmap [Internet]. 2018; Available from: https://github.com/raivokolde/pheatmap.

20. Gaspar LS, Santana MM, Henriques C, Pinto MM, Ribeiro-Rodrigues TM, Girao H, Nobre RJ, Pereira de Almeida L. Simple and Fast SEC-Based Protocol to Isolate Human Plasma-Derived Extracellular Vesicles for Transcriptional Research. Mol Ther Methods Clin Dev. 2020;18:723–37. doi: 10.1016/j.omtm.2020.07.012.

21. Welsh JA, Goberdhan DCI, O’Driscoll L, Buzas EI, Blenkiron C, Bussolati B, et al. Minimal information for studies of extracellular vesicles (MISEV2023): From basic to advanced approaches. J Extracell Vesicles. 2024;13(2):e12404. doi: 10.1002/jev2.12404.

22. Stephens M. False discovery rates: a new deal. Biostatistics. 2017;18(2):275–94. doi: 10.1093/biostatistics/kxw041.

23. Appierto V, Callari M, Cavadini E, Morelli D, Daidone MG, Tiberio P. A lipemia-independent NanoDrop((R))-based score to identify hemolysis in plasma and serum samples. Bioanalysis. 2014;6(9):1215–26. doi: 10.4155/bio.13.344.

24. ggplot2 [Internet]. Available from: https://github.com/tidyverse/ggplot2.

25. Rubio Gomez MA, Ibba M. Aminoacyl-tRNA synthetases. RNA. 2020;26(8):910–36. doi: 10.1261/rna.071720.119.

26. Sugimoto M, Kuo ML, Roussel MF, Sherr CJ. Nucleolar Arf tumor suppressor inhibits ribosomal RNA processing. Mol Cell. 2003;11(2):415–24. doi: 10.1016/s1097-2765(03)00057-1.

27. Rajan S, Chu Pham Dang H, Djambazian H, Zuzan H, Fedyshyn Y, Ketela T, Moffat J, Hudson TJ, Sladek R. Analysis of early C2C12 myogenesis identifies stably and differentially expressed transcriptional regulators whose knock-down inhibits myoblast differentiation. Physiol Genomics. 2012;44(2):183–97. doi: 10.1152/physiolgenomics.00093.2011.

28. Banerjee S, Chaturvedi R, Singh A, Kushwaha HR. Putting human Tid-1 in context: an insight into its role in the cell and in different disease states. Cell Commun Signal. 2022;20(1):109. doi: 10.1186/s12964-022-00912-5.

29. Arbour N, Tremblay P, Oth D. N-formyl-methionyl-leucyl-phenylalanine induces and modulates IL-1 and IL-6 in human PBMC. Cytokine. 1996;8(6):468–75. doi: 10.1006/cyto.1996.0063.

30. Bonavita CM, White TM, Francis J, Cardin RD. Heart Dysfunction Following Long-Term Murine Cytomegalovirus Infection: Fibrosis, Hypertrophy, and Tachycardia. Viral Immunol. 2020;33(3):237–45. doi: 10.1089/vim.2020.0007.

31. Wang Z, Xu H, Chen M, Lu Y, Zheng L, Ma L. CCL24/CCR3 axis plays a central role in angiotensin II-induced heart failure by stimulating M2 macrophage polarization and fibroblast activation. Cell Biol Toxicol. 2023;39(4):1413–31. doi: 10.1007/s10565-022-09767-5.

32. Ishida TK, Tojo T, Aoki T, Kobayashi N, Ohishi T, Watanabe T, Yamamoto T, Inoue J. TRAF5, a novel tumor necrosis factor receptor-associated factor family protein, mediates CD40 signaling. Proc Natl Acad Sci U S A. 1996;93(18):9437–42. doi: 10.1073/pnas.93.18.9437.

33. Gali VK, Balint E, Serbyn N, Frittmann O, Stutz F, Unk I. Translesion synthesis DNA polymerase eta exhibits a specific RNA extension activity and a transcription-associated function. Sci Rep. 2017;7(1):13055. doi: 10.1038/s41598-017-12915-1.

34. Frohlich ED, Susic D. Pressure overload. Heart Fail Clin. 2012;8(1):21-32. doi: 10.1016/j.hfc.2011.08.005.

35. Khan A, Pare E, Shah S. Peripartum Cardiomyopathy: a Review for the Clinician. Curr Treat Options Cardiovasc Med. 2018;20(11):91. doi: 10.1007/s11936-018-0690-3.

36. Bandla H, Dasgupta D, Mauer AS, Nozickova B, Kumar S, Hirsova P, Graham RP, Malhi H. Deletion of endoplasmic reticulum stress-responsive co-chaperone p58(IPK) protects mice from diet-induced steatohepatitis. Hepatol Res. 2018;48(6):479–94. doi: 10.1111/hepr.13052.

37. Hanafusa H, Torii S, Yasunaga T, Nishida E. Sprouty1 and Sprouty2 provide a control mechanism for the Ras/MAPK signalling pathway. Nat Cell Biol. 2002;4(11):850–8. doi: 10.1038/ncb867.

38. Organ SL, Tsao MS. An overview of the c-MET signaling pathway. Ther Adv Med Oncol. 2011;3(1 Suppl):S7-S19. doi: 10.1177/1758834011422556.

39. Oda K, Matsuoka Y, Funahashi A, Kitano H. A comprehensive pathway map of epidermal growth factor receptor signaling. Mol Syst Biol. 2005;1:2005 0010. doi: 10.1038/msb4100014.

40. Werner H. The IGF1 Signaling Pathway: From Basic Concepts to Therapeutic Opportunities. Int J Mol Sci. 2023;24(19). doi: 10.3390/ijms241914882.

41. Shkurnikov MY, Knyazev EN, Fomicheva KA, Mikhailenko DS, Nyushko KM, Saribekyan EK, Samatov TR, Alekseev BY. Analysis of Plasma microRNA Associated with Hemolysis. Bull Exp Biol Med. 2016;160(6):748–50. doi: 10.1007/s10517-016-3300-y.

42. Ayer A, Truby LK, Schroder JN, Casalinova S, Green CL, Bishawi MA, Bryner BS, Milano CA, Patel CB, Devore AD. Improved Outcomes in Severe Primary Graft Dysfunction After Heart Transplantation Following Donation After Circulatory Death Compared With Donation After Brain Death. J Card Fail. 2023;29(1):67–75. doi: 10.1016/j.cardfail.2022.10.429.

43. Kharawala A, Nagraj S, Seo J, Pargaonkar S, Uehara M, Goldstein DJ, Patel SR, Sims DB, Jorde UP. Donation After Circulatory Death Heart Transplant: Current State and Future Directions. Circ Heart Fail. 2024;17(7):e011678. doi: 10.1161/CIRCHEARTFAILURE.124.011678.

44. Iyer A, Gao L, Doyle A, Rao P, Jayewardene D, Wan B, et al. Increasing the tolerance of DCD hearts to warm ischemia by pharmacological postconditioning. Am J Transplant. 2014;14(8):1744–52. doi: 10.1111/ajt.12782.

45. White CW, Ali A, Hasanally D, Xiang B, Li Y, Mundt P, et al. A cardioprotective preservation strategy employing ex vivo heart perfusion facilitates successful transplant of donor hearts after cardiocirculatory death. J Heart Lung Transplant. 2013;32(7):734–43. doi: 10.1016/j.healun.2013.04.016.

46. White CW, Hasanally D, Mundt P, Li Y, Xiang B, Klein J, et al. A whole blood-based perfusate provides superior preservation of myocardial function during ex vivo heart perfusion. J Heart Lung Transplant. 2015;34(1):113–21. doi: 10.1016/j.healun.2014.09.021.

47. Saemann L, Wachter K, Georgevici AI, Pohl S, Hoorn F, Veres G, Korkmaz-Icoz S, Karck M, Simm A, Szabo G. Transcriptomic Changes in the Myocardium and Coronary Artery of Donation after Circulatory Death Hearts following Ex Vivo Machine Perfusion. Int J Mol Sci. 2024;25(2). doi: 10.3390/ijms25021261.

48. Schulze PC. Myocardial lipid accumulation and lipotoxicity in heart failure. J Lipid Res. 2009;50(11):2137–8. doi: 10.1194/jlr.R001115.

49. Zhang S, Hulver MW, McMillan RP, Cline MA, Gilbert ER. The pivotal role of pyruvate dehydrogenase kinases in metabolic flexibility. Nutr Metab (Lond). 2014;11(1):10. doi: 10.1186/1743-7075-11-10.

50. Drake KJ, Sidorov VY, McGuinness OP, Wasserman DH, Wikswo JP. Amino acids as metabolic substrates during cardiac ischemia. Exp Biol Med (Maywood). 2012;237(12):1369–78. doi: 10.1258/ebm.2012.012025.

51. Rodriguez GP, Song JB, Crouse GF. In vivo bypass of 8-oxodG. PLoS Genet. 2013;9(8):e1003682. doi: 10.1371/journal.pgen.1003682.

52. Wu L, Sowers JR, Zhang Y, Ren J. Targeting DNA damage response in cardiovascular diseases: from pathophysiology to therapeutic implications. Cardiovasc Res. 2023;119(3):691–709. doi: 10.1093/cvr/cvac080.

53. Crozier SJ, Zhang X, Wang J, Cheung J, Kimball SR, Jefferson LS. Activation of signaling pathways and regulatory mechanisms of mRNA translation following myocardial ischemia-reperfusion. J Appl Physiol (1985). 2006;101(2):576-82. doi: 10.1152/japplphysiol.01122.2005.

54. Chen X, Shi C, He M, Xiong S, Xia X. Endoplasmic reticulum stress: molecular mechanism and therapeutic targets. Signal Transduct Target Ther. 2023;8(1):352. doi: 10.1038/s41392-023-01570-w.

55. He X, Du T, Long T, Liao X, Dong Y, Huang ZP. Signaling cascades in the failing heart and emerging therapeutic strategies. Signal Transduct Target Ther. 2022;7(1):134. doi: 10.1038/s41392-022-00972-6.

56. Javadov S, Jang S, Agostini B. Crosstalk between mitogen-activated protein kinases and mitochondria in cardiac diseases: therapeutic perspectives. Pharmacol Ther. 2014;144(2):202–25. doi: 10.1016/j.pharmthera.2014.05.013.

57. Khavinson V, Linkova N, Dyatlova A, Kantemirova R, Kozlov K. Senescence-Associated Secretory Phenotype of Cardiovascular System Cells and Inflammaging: Perspectives of Peptide Regulation. Cells. 2022;12(1). doi: 10.3390/cells12010106.

58. Nog R, Aggarwal Gupta C, Panza JA. Role of MicroRNA in Heart Transplant. Cardiol Rev. 2022;30(5):253–7. doi: 10.1097/CRD.0000000000000393.

59. Landry P, Plante I, Ouellet DL, Perron MP, Rousseau G, Provost P. Existence of a microRNA pathway in anucleate platelets. Nat Struct Mol Biol. 2009;16(9):961–6. doi: 10.1038/nsmb.1651.

60. Sprenkle NT, Serezani CH, Pua HH. MicroRNAs in Macrophages: Regulators of Activation and Function. J Immunol. 2023;210(4):359–68. doi: 10.4049/jimmunol.2200467.

61. Heijnen HF, Schiel AE, Fijnheer R, Geuze HJ, Sixma JJ. Activated platelets release two types of membrane vesicles: microvesicles by surface shedding and exosomes derived from exocytosis of multivesicular bodies and alpha-granules. Blood. 1999;94(11):3791–9.

62. Boudreau LH, Duchez AC, Cloutier N, Soulet D, Martin N, Bollinger J, et al. Platelets release mitochondria serving as substrate for bactericidal group IIA-secreted phospholipase A2 to promote inflammation. Blood. 2014;124(14):2173–83. doi: 10.1182/blood-2014-05-573543.

63. Wacker J, Lucchinetti E, Jamnicki M, Aguirre J, Harter L, Keel M, Zaugg M. Delayed inhibition of agonist-induced granulocyte-platelet aggregation after low-dose sevoflurane inhalation in humans. Anesth Analg. 2008;106(6):1749–58. doi: 10.1213/ane.0b013e318172f9e9.

64. Yuki K, Bu W, Shimaoka M, Eckenhoff R. Volatile anesthetics, not intravenous anesthetic propofol bind to and attenuate the activation of platelet receptor integrin alphaIIbbeta3. PLoS One. 2013;8(4):e60415. doi: 10.1371/journal.pone.0060415.

65. Kirschner MB, Edelman JJ, Kao SC, Vallely MP, van Zandwijk N, Reid G. The Impact of Hemolysis on Cell-Free microRNA Biomarkers. Front Genet. 2013;4:94. doi: 10.3389/fgene.2013.00094.

66. Srikanth K, Park JE, Ji SY, Kim KH, Lee YK, Kumar H, et al. Genome-Wide Transcriptome and Metabolome Analyses Provide Novel Insights and Suggest a Sex-Specific Response to Heat Stress in Pigs. Genes (Basel). 2020;11(5). doi: 10.3390/genes11050540.

67. Yusifov A, Chhatre VE, Koplin EK, Wilson CE, Schmitt EE, Woulfe KC, Bruns DR. Transcriptomic analysis of cardiac gene expression across the life course in male and female mice. Physiol Rep. 2021;9(13):e14940. doi: 10.14814/phy2.14940.

